# Maturational indices of the cognitive control network are associated with inhibitory control in early childhood

**DOI:** 10.1101/2021.07.02.450852

**Authors:** Philipp Berger, Angela D. Friederici, Charlotte Grosse Wiesmann

## Abstract

Goal-directed behavior crucially relies on our capacity to suppress impulses and predominant behavioral responses. This ability, called inhibitory control, emerges in early childhood with marked improvements between 3 and 4 years. Here, we ask which brain structures are related to the emergence of this critical ability. Using a multimodal approach, we relate the pronounced behavioral improvements in different facets of 3-and 4-year-olds’ (N = 37, 20 female) inhibitory control to structural indices of maturation in the developing brain assessed with MRI. Our results show that cortical and subcortical structure of core regions in the adult cognitive control network, including the PFC, thalamus, and the inferior parietal cortices, are associated with early inhibitory functioning in preschool children. Probabilistic tractography revealed an association of frontoparietal (i.e., the superior longitudinal fascicle) and thalamocortical connections with early inhibitory control. Notably, these associations to brain structure were distinct for different facets of early inhibitory control, often referred to as motivational (‘hot’) and cognitive (‘cold’) inhibitory control. Our findings thus reveal the structural brain networks and connectivity related to the emergence of this core faculty of human cognition.

**Significance Statement:** The capacity to suppress impulses and behavioral responses is crucial for goal-directed behavior. This ability, called inhibitory control, develops between the ages of 3 and 4 years. The factors behind this developmental milestone have been debated intensely for decades, however, the brain structure that underlies the emergence of inhibitory control in early childhood is largely unknown. Here, we relate the pronounced behavioral improvements in inhibitory control between 3 and 4 years with structural brain markers of grey matter and white matter maturation. Using a multimodal approach that combines analyses of cortical surface structure, subcortical structures, and white matter connectivity, our results reveal the structural brain networks and connectivity related to this core faculty of human cognition.

## Introduction

The capacity to suppress impulses and behavioral responses is crucial for goal-directed behavior (Rothbart and Posner, 1985). This ability, called inhibitory control (IC), develops rapidly between the ages of 3 and 4 years (Petersen et al., 2016). The factors behind this developmental milestone have been debated for decades, however, the neurobiological correlates of the early development in IC are largely unknown. What are the brain structures related to this core faculty in human cognitive development?

In early childhood, the cerebral cortex undergoes complex developmental changes as a result of the interaction of microbiological processes (Raznahan et al., 2011; Walhovd et al., 2017). These maturational processes, observed in changes of cortical and subcortical grey matter (GM) and in the interconnectedness through white matter (WM) pathways, have each shown robust associations with cognitive development in childhood (Johansen-Berg, 2010; Walhovd et al., 2017). The interrelation of GM and WM structure with cognitive function, however, has only rarely been studied in the first years of life, given the difficulty to conduct high-resolution MRI in this age group (but see Cafiero et al., 2019). Therefore, to date, only little is known about the neural maturation processes underlying the emergence of IC in early childhood, particularly in the early preschool period from 3 years of age that marks a critical take-off in IC abilities.

In adults and older children, functional MRI studies have revealed that mature cognitive control is supported by a neural network of prefrontal and parietal cortices, along with subcortical structures (including the ventral striatum and thalamus), referred to as the superordinate Cognitive Control Network (sCCN) (Niendam et al., 2012; McKenna et al., 2017). In addition, connections between subcortical and the relevant cortical regions were shown to be involved in mature IC (Somerville and Casey, 2010; Halassa and Kastner, 2017). Studies in middle childhood and adolescence suggest some continuity of the functional involvement of the sCNN and PFC in cognitive control, at least from late childhood into adolescence (see McKenna et al., 2017; Fiske and Holmboe, 2019). For younger children, there is evidence from functional Near-Infrared Spectroscopy (fNIRS) studies for a similar frontoparietal network to be functionally involved between the age of 3 and 6 years (Mehnert et al., 2013; Buss and Spencer, 2018; Moriguchi et al., 2018; Moriguchi, 2021). These findings suggest that the brain-structural maturation of the sCCN may be associated with the emergence of early IC.

In behavioral studies on early IC development, different facets of inhibition have been described (Simpson and Carroll, 2019), including a distinction between inhibition in neutral and emotional task contexts, referred to as ‘cold’ and ‘hot’ IC (Zelazo and Carlson, 2012; Montroy et al., 2019). Neuroimaging research in adults and older children has largely focused on inhibition under motivationally neutral (‘cold’) task conditions. A standard ‘cold’ inhibition measure is the Go-NoGo task. In this task, children are required to perform a simple motor response when a stimulus is shown and to withhold this response when seeing another stimulus. Another line of the developmental research, in contrast, has focused on inhibition in emotional high-stake (‘hot’) situations. The standard task for this domain of IC is the Delay of Gratification tasks (Mischel et al., 1989). In this task, children are required to resist their impulse to choose an immediate reward in favor of a delayed larger reward. Although the distinction between ‘hot’ and ‘cold’ IC is widely recognized in the behavioral developmental literature, it is debated whether, despite the conceptual differences in task designs, these components share a common basis or address distinct and independent cognitive processes (Simpson and Carroll, 2019). From a neural perspective, studies in adults suggest that performance in the Delay of Gratification task might relate to a different functional network, including the orbitofrontal cortex (OFC) and frontostriatal connections (Casey et al., 2011). Studies in children, in turn, have reported functional recruitment of the lateral PFC (Moriguchi et al., 2018).

In the present study, we use high-resolution MRI to investigate how structural markers of brain maturation in GM and WM are related to early IC in the preschool period. In particular, we relate behavioral indices of different facets of early IC to structural properties of the PFC, OFC, and more broadly the sCCN, investigating cortical and subcortical structures in combination with WM connectivity in the critical age of 3-to 4 years when IC emerges.

## Materials and Methods

### Participants

MRI data and behavioral data of 37 typically developing 3-and 4-year-old children were analyzed for the present study (17 children aged 3.07–3.59 years, median=3.33, s.d.=0.18, 10 female; and 20 children aged 4.02–4.58 years, median=4.29, s.d.=0.18, 10 female). The sample size was estimated based on previous developmental MRI studies with approximately 20 children per age group, assuming a dropout rate of 10–20% due to motion artefacts. Thus, the behavioral assessment was conducted in a total sample of n = 60 children aged 3 and 4 years (Grosse Wiesmann et al., 2017a) from which we excluded children because they 1) did not participate in both IC tasks (n = 1), 2) did not participate in or aborted the MRI (n = 9), 3) showed incidental neurological findings (n = 1), 4) showed motion artifacts in the sMRI data (n = 11) (Grosse Wiesmann et al., 2020) or 5) dMRI data (Grosse Wiesmann et al., 2017b; see the respective sections on sMRI/dMRI data analysis for details). One additional child was excluded due to an MRI acquisition error. Parental informed consent was obtained for all children, and the study was approved by the Ethics Committee at the Faculty of Medicine of the University of Leipzig.

### Assessment of IC

The children performed two standard tests of ‘cold’ and ‘hot’ IC – a Go-NoGo task (Rakoczy, 2010) and a Delay of Gratification task (Mischel and Ebbesen, 1970), which have been described in detail in previous studies (Grosse Wiesmann et al., 2017b, 2017a). Briefly, in the Go-NoGo task, children were asked to perform actions a duck puppet asked them to do (for example, ‘Clap your hands!’), but not to do anything the nasty crocodile asked them to. A d-prime value was calculated with correct NoGo-trials as hits and incorrect Go-trials as false alarms (M=0.875, s.d.=0.178). One child had to be excluded from further analyses because they did not provide complete data on the task. For the Delay of Gratification task, children asked to wait in front of highly desirable object (gummy bears or chocolate bars) for 5 minutes to receive a bigger reward. One child had to be excluded from further analyses because they did not provide complete data on the task. The children’s mean waiting time was M=226□Js (s.d.=111s). To ensure comparability across task scores, we computed standardized scores of task performance in the Go-NoGo task and the Delay of Gratification task. Note that the data on these tasks were non-normally distributed because of the developmental breakthrough observed in the age range between 3 and 4 years. Therefore, we made sure that the statistical tests conducted were either non-parametric or that the data met the assumptions (i.e., normality of residuals).

### MRI data acquisition

MRI data were acquired on a 3-T Siemens scanner (Siemens MRT Trio series) using a 32-channel head coil. The acquisition protocols are described in detail in previous studies (Grosse Wiesmann et al., 2017b, 2020). High-resolution 3D T1-weighted MRI images were acquired using the MP2RAGE sequence (Marques et al., 2010) at 1.2 × 1 × 1 mm resolution (Grosse Wiesmann et al., 2020). Furthermore, dMRI data were acquired using the multiplexed echo planar imaging sequence (Feinberg et al., 2010) with a resolution of 1.9□mm isotropic (Grosse Wiesmann et al., 2017b). A field map was acquired directly after the dMRI scan.

### sMRI data analysis

#### Cortical surface-based analyses

To obtain measures of cortical thickness and surface area, we used the preprocessed brain images from a recent study using the current dataset (Grosse Wiesmann et al., 2020). In short, individual brain images were preprocessed in FreeSurfer 5.3.0 (http://surfer.nmr.mgh.harvard.edu) to reconstruct cortical surfaces and generate local estimates of cortical thickness and surface area following the standard surface-based stream (Fischl and Dale, 2000). Using a standardized workflow to detect motion artefacts (Backhausen et al., 2016), n = 11 children were excluded after initial screening, only including data of participants that had good to moderate MRI scans. Further, as recommended in the FreeSurfer pipeline (http://surfer.nmr.mgh.harvard.edu/fswiki/FsTutorial/TroubleshootingData), the outputs of skull stripping, white matter segmentation, and cortical and pial surfaces were inspected visually for errors and corrected manually when necessary. The automated FreeSurfer pipeline was rerun for the surfaces that contained errors and then reinspected (Backhausen et al., 2016). Surface area of the GM/WM boundary and cortical thickness, defined as the closest distance from the GM/WM boundary to the GM/CSF boundary, was calculated at each vertex. The resulting maps for cortical thickness and surface area were smoothed on the tessellated surfaces using a 10-mm FWHM Gaussian kernel. A common group template was created from the individual T1-weighted images of all children included in the analysis using ANTs (Avants et al., 2008). The individual cortical surfaces were registered to the common group template to allow for an accurate matching of local cortical thickness and surface area measures across participants.

#### Subcortical volume analysis

Individual brain images were processed in FreeSurfer 7.0.0 (http://surfer.nmr.mgh.harvard.edu) to reconstruct subcortical volumes following the standard volume-based processing stream in FreeSurfer. The volume-based FreeSurfer processing stream conducts segmentation of several subcortical structures including the thalamus and striatum. Based on our hypotheses, we restricted subsequent analyses to the bilateral thalamus and structures of the striatum, including the bilateral putamen, bilateral caudate nucleus and bilateral nucleus accumbens. Quality control assessments of the thalamic and striatal reconstructions was conducted by two independent raters, yielding an interrater reliability of M_ICC_=.70 (s.d.= .08). If the two raters agreed on a structure to show failures in segmentation this structure was excluded from the statistical analyses for the individual participant. This procedure resulted in the exclusion of the following number of participants per structure: N = 1 for the left caudate nucleus, N = 2 for the right caudate nucleus, N = 2 for the left putamen, N = 2 for the right putamen, N = 5 for the left nucleus accumbens, N = 6 for right nucleus accumbens, N = 1 for the left thalamus, and N = 1 for right thalamus.

#### Statistical analysis

The relation of cortical thickness and surface area, respectively, with our main variables (‘cold’ and ‘hot’ IC) were estimated in general linear models (GLM) using the tool mri_glmfit implemented in FreeSurfer. In the GLMs, we controlled for children’s chronological age, and gender. Multiple comparison correction was applied with a clusterwise correction using the FreeSurfer tool mri_glmfit-sim, specifying a cluster-forming threshold of P < 0.005, clusterwise threshold of P < 0.05 (Greve and Fischl, 2018), positive relation with surface area, bidirectional relation with cortical thickness, and additional correction for the analyses on two hemispheres (Grosse Wiesmann et al., 2020). For the clusterwise correction, a Monte Carlo simulation with 10,000 iterations was precomputed on the group template. Given our hypotheses for neural networks associated with ‘cold’ and ‘hot’ IC, respectively, we computed additional analyses using small-volume correction. In particular, we used a binarized map of the sCCN, derived from a meta-analysis on ‘cold’ inhibitory functioning in adults (Niendam et al., 2012). For this, we registered the original meta-analysis maps from the MNI space to our group template with the ANTs script WarpImageMultiTransform (http://stnava.github.io/ANTs/) and then projected them on the surface using the FreeSurfer tool mri_vol2surf (see Figure 1A). Furthermore, based on our specific hypotheses for PFC and OFC, we selected these as regions of interest based on the Desikan-Killiany parcellation (Desikan et al., 2006). The linear models for the relations of cortical thickness and surface area with ‘cold’ and ‘hot’ IC, respectively (as well as the analyses that included the covariates described above) were computed within the obtained masks with mri_glmfit as before. We checked for the normality of residuals using Kolmogorov-Smirnov tests.

**Figure 1.**
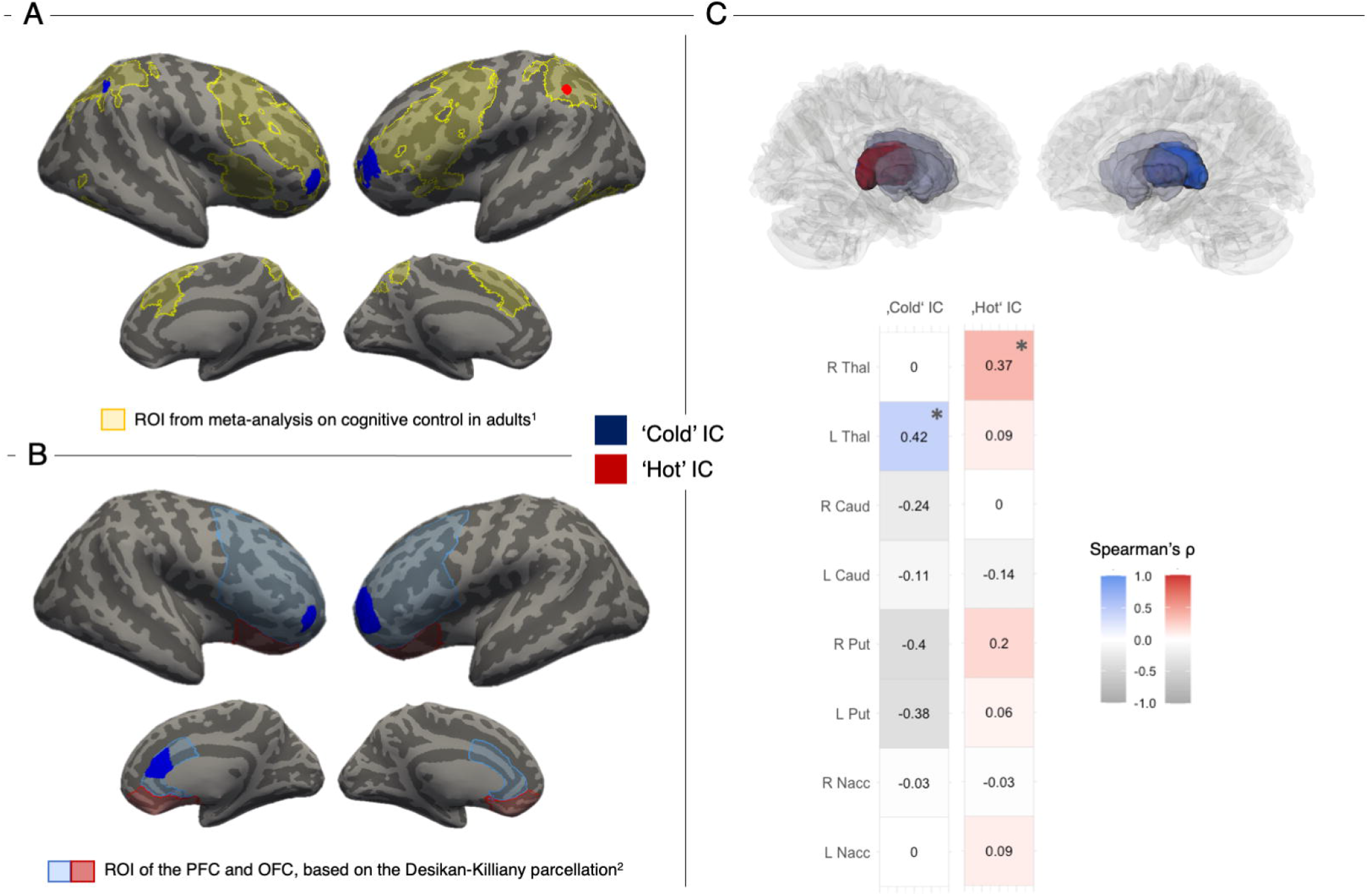
Relation of ‘cold’ (blue) and ‘hot’ (red) inhibitory control (IC) performance with cortical surface area and subcortical volume. (A) Small volume corrected linear relation of ‘cold’ IC performance (blue) with surface area in bilateral rostral frontal cortex (BA10) and right inferior parietal lobe and ‘hot’ IC performance (red) with cortical surface area in the left supramarginal gyrus within the regions of the superordinate Cognitive Control Network (sCCN; light yellow). (B) Small-volume corrected linear relation of ‘cold’ IC performance with surface area in the caudal ACC within the prefrontal cortex (PFC) mask (light blue) and medial orbitofrontal cortex (OFC) within the OFC mask (light red). All effects on the cortical surface are independent of children’s age and gender, and cluster-size corrected at P < 0.05. (C) Results of non-parametric partial correlation analyses of ‘cold’ and ‘hot’ IC performance with volume of subcortical structures, color-coded for strength of positive association. A significant positive correlation of ‘cold’ IC (blue) with volume of the left thalamus, and ‘hot’ IC (red) with the right thalamus was revealed. These relations were independent of children’s chronological age and gender. Abbreviations: Thal = Thalamus, Caud = Caudate Nucleus, Put = Putamen, Nacc = Nucleus accumbens.

The relation of subcortical volume of the thalamus and striatum with IC scores was assessed with non-parametric partial correlation analysis, using the ‘ppcor’ package (Kim, 2015) in R (R Core Team, 2019). In the analysis, we used a one-sided (positive) test to indicate significant correlations, while controlling for children’s chronological age, gender and eTIV, additionally correcting for the number of structures via false discovery rate (Benjamini and Hochberg, 1995).

### dMRI data analysis

#### dMRI data processing

To obtain measures of WM connectivity, we used the preprocessed dMRI data and analysis pipeline from another study using the current dataset (Grosse Wiesmann et al., 2017b). To ensure reliable data quality, children were excluded if more than 10 out of 60 acquired directions in the dMRI data set were corrupted. Directions were removed due to intensity dropout caused by head motion (Schreiber et al., 2014) or due to artefacts detected in a visual inspection (Tournier et al., 2011; Soares et al., 2013). After removing motion artefacts manually, motion itself was corrected for by rigidly aligning all volumes to the last one without diffusion weighting (b0) using flirt from the FSL software package (Jenkinson et al., 2002). Subsequently, the motion corrected dMRI data were rigidly aligned to the anatomical image, which again had been rigidly aligned to the Montreal Neurological Institute (MNI) standard space and was interpolated to 1□mm isotropic voxel space. Distortions were corrected using the corresponding field map.

#### Seed region derivation

To see within which tracts the significant clusters from the sMRI analyses were located, these regions were taken as seeds for probabilistic tractography. For this, we projected the significant regions derived in the cortical surface-based and volume-based analysis on the GM/WM boundary in the individual subject’s brain using the FreeSurfer tool label2label. After transforming the label files to volume, we rigidly aligned them to the diffusion space using flirt from the FSL software package (Jenkinson et al., 2002). In addition, a supplementary analysis was performed focusing only on frontostriatal connectivity. As a seed region for probabilistic tractography, we used a mask of the OFC, based on the Desikan-Killiany parcellation in FreeSurfer. Further, we defined an inclusion region of interest combining FreeSurfer segmentations of bilateral caudate nucleus, putamen and nucleus accumbens, yielding that only tracks that enter the inclusion region of interest will be produced.

#### Statistical analysis

Tractography was run with MRtrix (Tournier et al., 2012) using Constrained Spherical Deconvolution as a local model (Tournier et al., 2004) with the default parameters, as in Grosse Wiesmann et al. (2017b). This procedure resulted in streamline density maps for each seed region and subject, in addition producing a map including only frontostriatal connections. To ensure that the identified correlations were not outlier-driven, we masked the streamline density maps of the individual subjects with a common group mask imposing that at least half the subjects have nonzero values in every voxel. The individual subjects’ masked streamline density maps were then correlated with IC scores using GLMs with non-paremtric permutation tests in FSL randomise (Winkler et al., 2014), while controlling for the mean volume in the seed region of the tractography. This was done in order to ensure that correlations with streamline density were not driven by the correlation of the IC score and GM structure found in the sMRI analysis. In addition, we controlled for age and gender by including them as covariates in the linear model. Reported clusters in the tract volumes were significant at P<0.005 at voxel-level and exceeded a cluster size significant at P<0.05, in addition taking to account the number of streamline density maps according to Bonferroni correction. We localized and named the clusters and tracts based on the MRI Atlas of Human White Matter (Oishi et al., 2011).

## Results

### Behavioral results

A behavioral assessment of ‘hot’ and ‘cold’ inhibitory functioning and codeveloping cognitive functions was conducted in 60 children aged 3 to 4 years (Grosse Wiesmann et al., 2017a), from which a subsample of 37 children with usable MRI data was analyzed with respect to cortical brain structure (for details, see Methods). The children performed two standard tasks for ‘cold’ and ‘hot’ inhibitory functioning – a Go-NoGo task and a Delay of Gratification task (details see Methods). Performance in the ‘cold’ IC task showed a significant correlation with age (Spearman’s ρ = .45, p < .001), with 4-year-olds (M=0.94, s.d.= .10) performing significantly better than 3-year-olds (M=0.74, s.d.=0.31; Mann-Whitney U-test, U = 202.50, p < 0.001, η^2^ = 0.20). Similarly, performance in the ‘hot’ IC task showed a significant correlation with age (Spearman’s ρ = .38, p = .003), with 4-year-olds (M=246.13 s, s.d.=94.62) performing significantly better than 3-year-olds (M=183.96 s, s.d.=119.35; Mann-Whitney U-test, U = 296.50, p = 0.039, η^2^ = 0.07). No significant correlation between ‘hot’ and ‘cold’ IC performance was found (Spearman’s ρ = .17, p = .224).

### Analysis of cortical brain structure

To test whether IC performance was related to cortical brain structure, we reconstructed cortical surface area and thickness from high-resolution anatomical MRI in the same children that participated in the behavioral task battery.

*‘Cold’ IC and cortical brain structure (Go-NoGo task)*

On the whole brain, we observed a positive correlation of children’s ‘cold’ IC performance with their cortical surface area in the left rostral frontal cortex (RFC, BA 10; r =.43, cluster-wise p = 0.005), inferior temporal gyrus (r =.54, cluster-wise p = 0.010), precentral gyrus (r =.45, cluster-wise p = 0.022), and right posterior cingulate cortex (Table 1; r =.48, cluster-wise p = 0.009). Given our a-priori hypotheses based on the brain regions relevant for ‘cold’ IC in adults, we additionally computed small-volume corrections within the regions of the sCCN (as reported in Niendam et al., 2012) and the PFC. In addition to an effect in the left RFC (r =.34, cluster-wise p < 0.001), this showed a positive correlation of children’s ‘cold’ IC performance with their cortical surface area in the right RFC (r =.36, cluster-wise p = 0.001), right inferior parietal lobe (IPL; r =.39, cluster-wise p = 0.015), and in the caudal ACC (r =.46, cluster-wise p = 0.001; Table 1, Fig.1A and 1B). All effects remained significant when controlling for chronological age and gender. There were no significant effects for cortical thickness.

**Table 1.** MNI coordinates, effect size, exact significance, and cluster size of significant relations between IC performance and cortical surface area.

| Anatomical region | Peak voxel coordinate<br>in MNI 305 space (X,<br>Y, Z) |  |  | Correlation<br>in peak<br>voxel | Cluster-wise<br>P value | Cluster-size<br>(in mm<sup>2</sup>) |
| --- | --- | --- | --- | --- | --- | --- |
| Whole-brain analysis |  |  |  |  |  |  |
| ‘Cold’ IC |  |  |  |  |  |  |
| L RFC | -35.4 | 57.1 | -6.2 | 0.431 | 0.00459 | 687.43 |
| L inf. temporal lobe | -63.3 | -43.2 | -20.2 | 0.544 | 0.00978 | 628.70 |
| L precentral gyrus* | -28.6 | -22.0 | 59.0 | 0.454 | 0.02247 | 550.10 |
| R PCC* | 13.6 | -23.9 | 35.9 | 0.480 | 0.00938 | 614.13 |
| Small-volume correction: sCCN |  |  |  |  |  |  |
| ‘Cold’ IC |  |  |  |  |  |  |
| L RFC* | -33.1 | 56.2 | -2.1 | 0.341 | 0.00020 | 376.86 |
| R RFC* | 32.2 | 53.8 | -4.7 | 0.358 | 0.00060 | 209.65 |
| R IPL* | 44.0 | -61.2 | 45.2 | 0.390 | 0.01460 | 136.17 |
| ‘Hot’ IC |  |  |  |  |  |  |
| L SMG* | -51.5 | -49.8 | 59.0 | 0.216 | 0.01720 | 113.21 |
| Small-volume correction: PFC |  |  |  |  |  |  |
| ‘Cold’ IC |  |  |  |  |  |  |
| L RFC* | -35.4 | 57.1 | -6.2 | 0.420 | 0.00010 | 687.43 |
| R RFC* | 38.0 | 58.8 | -7.1 | 0.387 | 0.03790 | 217.05 |
| R caudal ACC | 8.7 | 40.0 | 9.7 | 0.459 | 0.00120 | 374.78 |

Note. All effects are controlled for children’s chronological age and gender. \* Regions marked with an asterisk were independent from the other inhibitory control (IC) domain. Abbreviations: ACC = Anterior Cingulate Cortex, IPL = Inferior Parietal Lobe, OFC = Orbitofrontal Cortex, PCC = Posterior Cingulate Cortex, RFC = Rostral Frontal Cortex, SMG = Supramarginal Gyrus.

*‘Hot’ IC and cortical brain structure (Delay of Gratification task)*

To identify cortical brain structure associated with ‘hot’ IC development, we computed the linear relation of children’s performance in the Delay of Gratification task with cortical surface area and thickness on the whole cortical surface. No significant correlation was found with either surface area or cortical thickness on the whole brain level. In a next step, we then tested whether children’s ‘hot’ IC performance was related to cortical structure in our a-priori defined regions-of-interest. Using small-volume correction within the regions of the sCCN, PFC and OFC, we found a significant positive relation of children’s ‘hot’ IC performance with their cortical surface area in the left supramarginal gyrus (r =.22, cluster-wise p = 0.017; Table 1, Fig1B), and no significant relations in the PFC or OFC. Again, the effect remained significant when controlling for chronological age and gender.

*Dissociation of ‘cold’ and ‘hot’ IC*

To test for the independence of the effects observed for ‘cold’ and ‘hot’ IC, we additionally controlled for performance in the other task, respectively. This showed that the regions of significant relation with ‘cold’ IC performance in bilateral RFC and right IPL were independent of ‘hot’ IC performance. Similarly, the ‘hot’ IC effect observed in the supramarginal gyrus was independent of ‘cold’ IC performance. Only the effect in caudal ACC did not remain significant when controlling for ‘hot’ IC performance.

### Analysis of subcortical brain structure

To test whether IC performance was related to subcortical brain structure, we performed non-parametric partial correlational analysis, relating IC performance with the reconstructed volume of the thalamus and striatal areas, while controlling for age, gender and estimated total intracranial volume (eTIV). We observed a positive correlation between children’s ‘cold’ IC performance and the volume of the left thalamus (Spearman’s ρ = .42, p = .013, Figure 1C), whereas ‘hot’ IC performance was significantly related to the volume of the right thalamus (Spearman’s ρ = .37, p = .013, Figure 1C). The effect of ‘cold’ IC in the left thalamus remained significant after controlling for ‘hot’ IC performance. No significant correlation was found with the volume of striatal regions for ‘hot’ and ‘cold’ IC.

### Analysis of white matter connectivity

To investigate the structural connectivity of cortical and subcortical regions that were associated with ‘hot’ and ‘cold’ IC, respectively, we performed probabilistic tractography seeding in those regions. This analysis yielded a bilateral frontoparietal structural network connected to the cortical seed regions whose cortical structure had shown a linear relation to ‘cold’ and ‘hot’ IC performance (see Figure 2AB). In particular, seeding from clusters in the RFC and IPL clusters, we observed white matter structures such as the superior longitudinal fasciculus (SLF) and inferior fronto-occipital fasciculus, connecting the prefrontal cortex with temporoparietal regions. Through seeding in the caudal ACC, we observed structures such as the cingulum and corpus callosum. Further, seeding in the thalamus, we identified thalamocortical structural connections in both hemispheres (Figure 2C). Additionally, we obtained frontostriatal connections through seeding in the OFC and using inclusion regions of interest in the striatum (see Figure 3).

**Figure 2.**
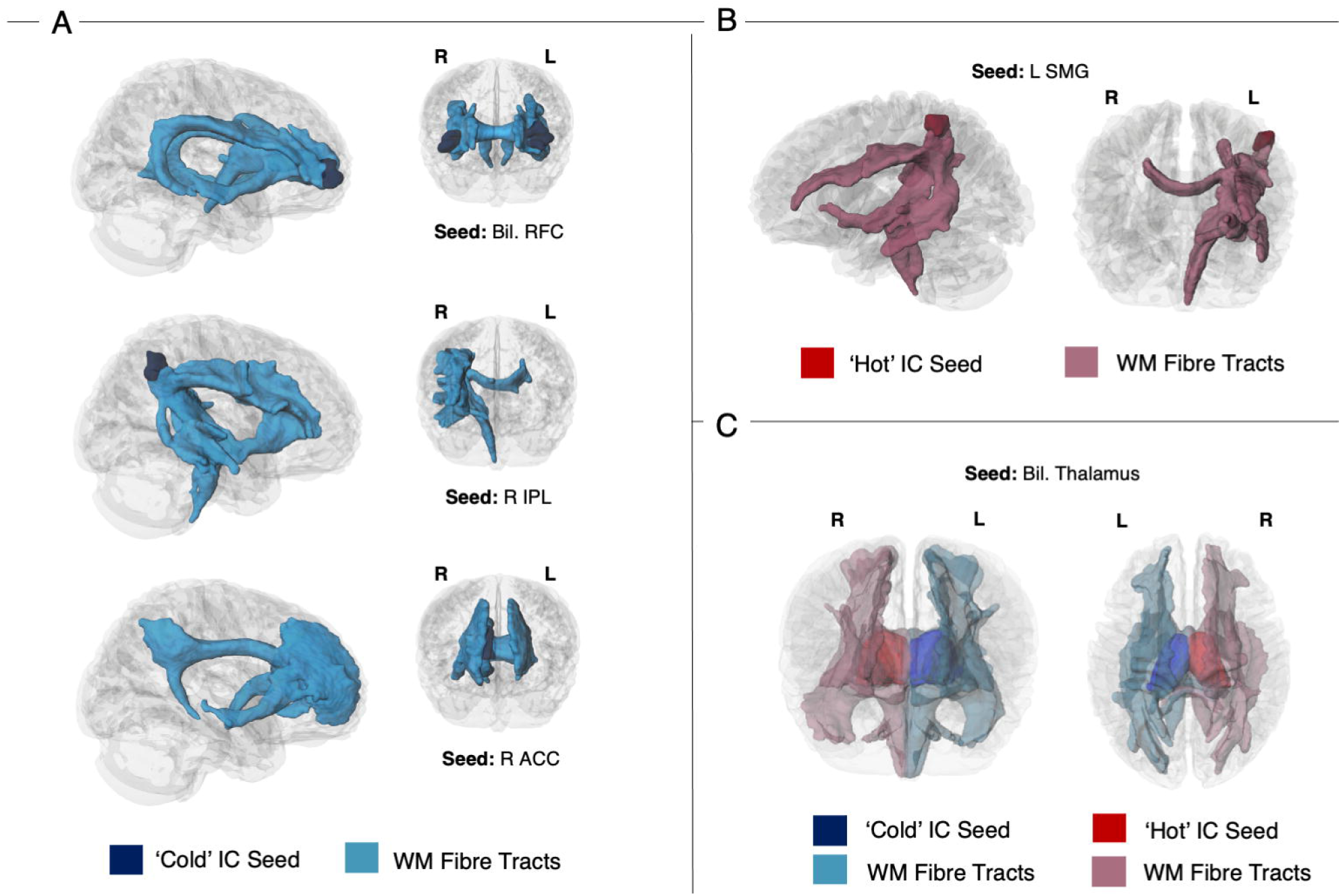
Streamline density maps resulting from probabilistic tractography seeded in brain regions showing a linear relation with ‘cold’ (blue) and ‘hot’ inhibitory control (IC) performance (red) on the cortical surface (A and B, respectively) and subcortical volume (bilateral thalamus, C). Abbreviations: ACC = Anterior Cingulate Cortex, IC = Inhibitory Control, IPL = Inferior Parietal Lobe, OFC = Orbitofrontal Cortex, RFC = Rostral Frontal Cortex, SMG = Supramarginal Gyrus, WM = White Matter.

**Figure 3.**
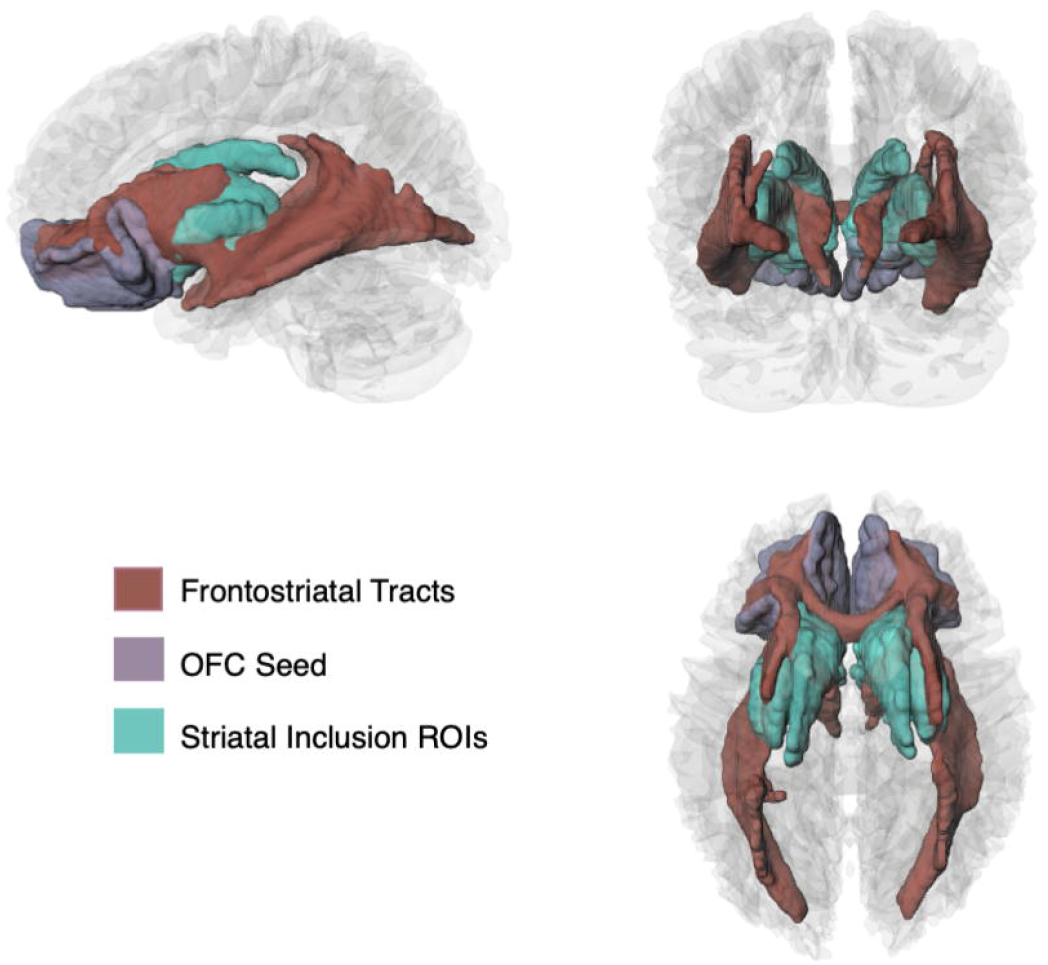
Streamline density map resulting from probabilistic tractography seeded in the orbitofrontal cortex (OFC; grey), using inclusion regions of interest (ROIs) in the striatum (green). The tractography revealed frontostriatal connections, along with the bilateral inferior fronto-occipital fasciculus (red).

Next, we wanted to specify how and where structural connectivity in these tracts was related to children’s IC abilities. To this end, we correlated streamline densities obtained from tractography with the children’s IC performance, while controlling for children’s age, gender and surface area/volume of the seed region (see Methods for details). This showed a significant correlation of ‘cold’ IC performance with streamline density in the DLPFC indexing higher connectivity to this region from the seed in the IPL along the right SLF (r = .47, p = 0.002) (see Figure 4B, Table 2). Furthermore, there were significant correlations of ‘cold’ inhibitory functioning with streamline densities along the major forceps (cluster 1: r = .47, p = 0.023; cluster 2: r =.46, p < 0.001) and in the left corticospinal tract (r = .44, p = 0.007), which connects the left thalamus with the motor cortex (see Figure 4A, Table 2), independently of ‘hot’ IC performance. For ‘hot’ IC, we obtained a significant correlation with streamline density of the right superior and posterior thalamic radiation indexing higher connectivity from the seed in the right thalamus to the frontoparietal and occipital cortex (r = .42, p = 0.001; see Figure 4C, Table 2). No significant correlation was obtained for ‘hot’ IC performance with streamline density of frontostriatal connections.

**Table 2.** MNI coordinates, effect size, exact significance, and cluster size of significant relations between IC performance and streamline density in WM tracts.

| Seed Region | Region of Cluster | Tractography WM tracts | MNI coordinates Centre of Gravity |  |  | Correlation coefficient | P value | Cluster-size (in voxels) |
| --- | --- | --- | --- | --- | --- | --- | --- | --- |
| ‘Cold’ IC |  |  |  |  |  |  |  |  |
| R IPL | R DLPFC | R SLF | 20.8 | 13.9 | 36.2 | 0.47 | 0.002 | 127 |
| L Thal | CC | Major Forceps* | 4.9 | -31.6 | 19.1 | 0.47 | 0.0230 | 123 |
| L Thal | CC | Major Forceps | -13.3 | -41.1 | 22.6 | 0.46 | < 0.001 | 83 |
| L Thal | L PoCG | L Corticospinal Tract* | -14 | -26.6 | 49.7 | 0.44 | 0.007 | 60 |
| ‘Hot’ IC |  |  |  |  |  |  |  |  |
| R Thal | R Thal | R Thalamic Radiation | 10.9 | -27.9 | 19.7 | 0.42 | 0.001 | 64 |

Note. All effects are controlled for children’s chronological age and gender, as well as surface area/volume of the respective seed region. Regions independent from the other IC domain are marked with an asterisk. P-values surviving Bonferroni correction for the number of tracts are marked with a plus sign. Abbreviations: DLPFC = Dorsolateral PFC, CC = Corpus Callosum, PoCG = Postcentral Gyrus, IPL = Inferior Parietal Lobe, SLF = Superior Longitudinal Fasciculus, WM = White Matter

**Figure 4.**
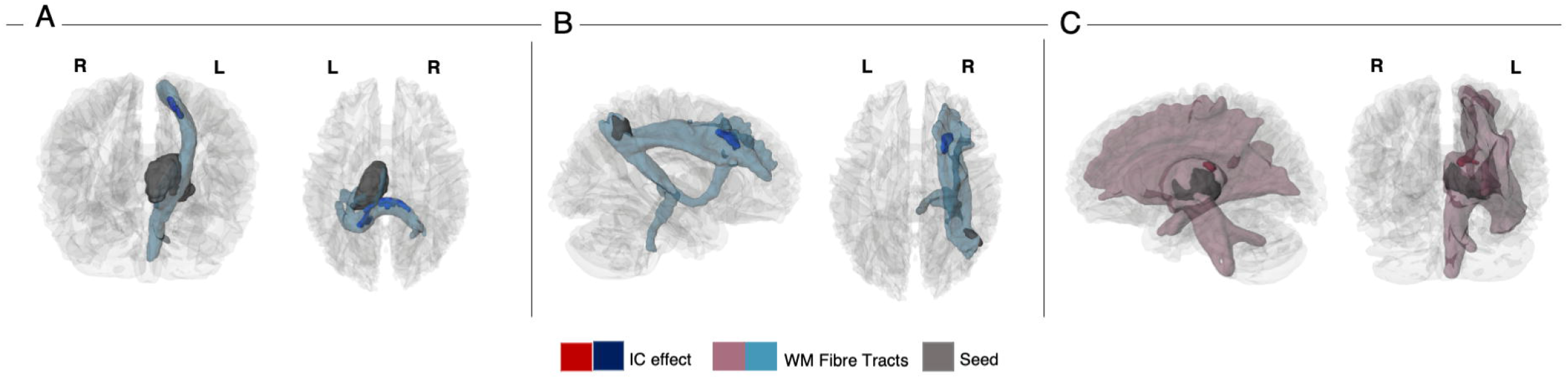
Correlation of streamline density with ‘cold’ (blue) and ‘hot’ inhibitory control (IC) performance (red). Results show a significant correlation of ‘cold’ IC with streamline density in the left corticospinal tract and forceps major (A), and in the right SLF (B). ‘Hot’ IC performance was correlated with streamline density of the right superior thalamic radiation (C). For visualization, white matter (WM) visitation maps (light red, light blue) are restricted to streamlines that pass through the significant cluster. The effects were independent of children’s age and gender, as well as surface area/volume of the seed region

## Discussion

IC constitutes a key prerequisite for humans to engage in goal-directed and adaptive behavior. A milestone of this ability is reached in the preschool period, between the ages of 3 and 4 years. What are the brain structures related to this core faculty in human cognitive development? To address this question, we studied structural brain markers of GM and WM maturation in relation to different facets of early IC cross-sectionally at the critical age of 3 to 4 years. We found that the behavioral breakthrough in IC was associated with structural indices of maturation in the cognitive control network, previously identified in adults (Niendam et al., 2012) and older children (McKenna et al., 2017). Distinct components of the cognitive control network were related to the different facets of IC: A standard cognitive (‘cold’) IC task was related to structure of the PFC, parietal lobe, and left thalamus, as well as frontoparietal and thalamocortical connections. A standard emotional (‘hot’) IC task, in contrast, was related to structural indices of maturation of the supramarginal gyrus, right thalamus, and distinct thalamocortical connections. These findings shed light on the brain structures associated with the emergence of IC, a central capacity for adaptive human behavior. Moreover, the independent brain networks involved in the different facets of IC support a dissociation of the structural networks underlying different task types thought to tap into ‘cold’ and ‘hot’ IC.

The emergence of ‘cold’ IC in early childhood was associated with structural indices of maturation in core regions in the sCCN. More specifically, surface area in prefrontal (i.e., ACC, bilateral RFC) and parietal cortices (i.e., right inferior parietal lobe), as well as volume of the left thalamus was related to the marked developmental improvement observed on classic ‘cold’ IC tasks between 3 and 4 years. A similar brain network has previously been found to be functionally involved in mature ‘cold’ IC tasks in adults and late childhood (Niendam et al., 2012; McKenna et al., 2017). These findings support that the breakthrough observed in ‘cold’ IC measures in preschool children (Petersen et al., 2016), such as in variants of the Go-NoGo task, reflects a development towards mature cognitive control. Structural connectivity of the above regions implicated in the ‘cold’ IC task revealed how these regions are integrated in structural networks. This analysis yielded both frontoparietal and thalamocortical pathways, connecting important hubs of the sCCN. In particular, performance in the Go-NoGo task was related to connectivity of the SLF connecting the DLPFC with the inferior parietal lobe, the posterior corpus callosum (i.e., major forceps) connecting the two hemispheres, and connections between the left thalamus and motor cortex through the left corticospinal tract. These results suggest that, in addition to brain structure in IC processing regions, the extent to which these regions are connected to each other and to the thalamus via association, projection and commissural fibres is relevant for the emergence of mature IC. Our findings highlight the importance of white matter fiber connections that enable communication between core regions of cognitive control in prefrontal and parietal cortices and the thalamus. In particular, the SLF has been shown to provide the prefrontal cortex with information on visual space and visuo-motor function processed in the inferior parietal cortices, which may contribute to the regulation of spatial attention (Mesulam, 1981; Posner et al., 1984; Bisley and Goldberg, 2003). The importance of the SLF for cognitive control is further supported by research on pathological conditions, showing that structural connectivity of the SLF is altered in disorders such as Obsessive-Compulsive Disorder (Gan et al., 2017) and Attention Deficit/Hyperactivity Disorder (ADHD; Hamilton et al., 2008; Gehricke et al., 2017). Specifically, research in ADHD has revealed that dysfunctional coupling in the SLF is associated with saccadic abnormalities (Fried et al., 2014; Matsuo et al., 2015) and increased response time variability (Wolfers et al., 2015). The corticospinal tract, in contrast, has functionally been linked to the control of voluntary movements (Kolb and Whishaw, 2009), and alterations of the corticospinal tract have been associated with the hyperactivity syndrome in ADHD (Bu et al., 2020). Our results, therefore, suggest that brain structures subserving different functional components of cognitive control, including attention regulation and motor control, are associated with young children’s IC in neutral contexts.

In contrast to IC in neutral contexts, inhibition in an emotionally and motivationally high-stake situation, as assessed with the standard ‘hot’ IC task, namely the Delay of Gratification task (Mischel et al., 1989), was related to structural indices of maturation in brain regions of the sCCN that were distinct from those observed for the Go-NoGo task. These included the supramarginal gyrus and right thalamus. This distinction was confirmed by a double dissociation in that the main reported effects were independent of performance in the respective other IC domain. This was also supported by a distinct structural network for the Delay of Gratification task, which was associated with the connectivity of the right thalamus with frontoparietal and occipital cortices via the superior and posterior thalamic radiation. Through ascending and descending tracts from the cerebral cortex the thalamic radiations integrate information throughout the brain, and are known to play a role in cognitive control and attention (Chaddock-Heyman et al., 2013; Stave et al., 2017; Brandes-Aitken et al., 2019).

In adults, the OFC and frontostriatal connections had previously been found to be functionally involved in IC in motivationally or emotionally-laden situations that require the inhibition of direct incentive needs to achieve a higher-value reward (Casey et al., 2011; Achterberg et al., 2016). In preschoolers, however, maturational indices of these structures were not significantly associated with performance in the Delay of Gratification task. This was further confirmed by an additional analysis focusing only on frontostriatal structures, but yielding no significant correlation of streamline density obtained from the tractography with children’s Delay of Gratification. Our data thus suggest that Delay of Gratification observed in the critical age range between 3 and 4 years is related to distinct components of the sCCN, rather than to a network specific for mature ‘hot’ IC. Indeed, besides the representation of incentive value emphasized in adults, ‘hot’ IC is thought to require the cognitive control of approach-avoidance tendencies (Zelazo and Carlson, 2020). For example, to master the Delay of Gratification task, children must not only represent and navigate between the values of immediate and delayed rewards, but also be able to suppress their motor responses appropriately and, if necessary, reorient their attention in a different direction to achieve the goal. Our finding that structural indices of the supramarginal gyrus, a region implicated in attentional control (Corbetta et al., 2008), and cortical connections of the thalamus to parietal regions and the motor cortex are related to early ‘hot’ IC suggests that controlling approach-avoidance tendencies may be critical for the emergence of this ability in early childhood. Whether and from which age the maturation of frontostriatal connections (Casey et al., 2011; Achterberg et al., 2016) may be relevant to the early development of ‘hot’ IC abilities remains a question for future investigation.

The observed differences in the structural networks related to ‘cold’ and ‘hot’ IC, respectively, are in line with evidence showing low behavioral associations of IC in ‘hot’ and cold’ task situations but synchronous development of the two capacities (Willoughby et al., 2011; Zelazo and Carlson, 2012; Montroy et al., 2019). Moreover, this is consistent with a dissociation of these abilities observed in clinical populations (Bechara, 2004; Eslinger et al., 2004) and developmental disorders, such as ADHD, which has been found to differentially impair ‘hot’ and ‘cold’ IC performances (Hobson et al., 2011; Antonini et al., 2015). On the neural level, our brain-structural results complement recent findings from an fNIRS study by Moriguchi (2021), showing that PFC activation during ‘cold’ IC tasks was generally stronger and uncorrelated to activation in the classical ‘hot’ Delay of Gratification task. Taken together, these findings corroborate that the neural basis of ‘hot’ and ‘cold’ IC may differ in early childhood.

Notably, however, in light of the increasingly consistent findings supporting a dissociation of ‘hot’ and ‘cold’ IC abilities on the behavioral, neural and clinical level, it is crucial to consider the inconsistencies in the way ‘hot’ and ‘cold’ IC are defined and studied. In particular, due to their different research backgrounds, standard ‘cold’ and ‘hot’ inhibition tasks, such as the Go-NoGo task and the Delay of Gratification task, do not only differ in the level of emotion or motivation included. They also show fundamental differences on many other factors in the task design (Simpson and Carroll, 2019). Different possibilities to conceptualize the differences between these tasks have been suggested in the developmental literature, such as, ‘conflict’ versus ‘delay’-based inhibition (Carlson and Moses, 2001), or, ‘strength’ versus ‘endurance’-based inhibition (Simpson and Carroll, 2019). Against this background, the question arises whether the observed dissociation of ‘hot’ and ‘cold’ IC may reflect other differences captured by these tasks. Future research could use several tasks for the IC domains in order to weaken the influence of task designs. However, in order to fully address this question in future research, it might be necessary to go beyond the traditional ‘hot-cold’ distinction, which relies on differences in the operationalization of ‘hot’ and ‘cold’ executive functions (Zelazo and Carlson, 2012). Instead, the influence of emotion and motivation on inhibitory processing may be studied as a factor within the same task. This would ensure that task requirements do not differ besides the level of emotion included. Previous research in adults found that the ventral ACC modulated the influence of emotion on IC with such task designs (Kanske and Kotz, 2011; Kanske, 2012). Recently, similar tasks have been developed for preschool children (Berger and Grosse Wiesmann, 2021) and older children (Zinchenko et al., 2019). These tasks could be used to study associated brain maturation with a similar approach as the one used in the present study.

In sum, the present findings suggest that cortical and subcortical structure of core regions in the adult cognitive control network are related to early inhibitory functioning in preschool children. Further, our findings highlight the importance of frontoparietal (i.e., the superior longitudinal fascicle) and thalamocortical connections for early inhibitory control. Notably, however, the observed associations to brain structure were distinct for different facets of early inhibitory control, previously suggested in the behavioral literature (Zelazo and Carlson, 2012; Simpson and Carroll, 2019).

## Acknowledgements

We would like to thank Julia Werner, Anne Grigutsch, Lisa Uhlich, and Alina Kowald for their help with correcting segmentation and surface reconstruction in FreeSurfer, and Hung Nguyen Trong and Christiane Attig for their help with data acquisition and organization.

## Author Contributions

C.G.W. and A.D.F. conceived the research; C.G.W. designed the tasks and conducted the research; P.B. and C.G.W. analyzed the data; P.B. wrote the paper; and C.G.W. and A.D.F. edited the paper.

## Notes

### Competing Interest Statement

The authors have declared no competing interest.

### Summary of Updates

- Revised introduction and discussion - Added information to methods - Added Figure to show frontostriatal connections - Added information on effect sizes

